# Rudhira-mediated microtubule stability controls TGFβ signaling during mouse vascular development

**DOI:** 10.1101/2024.04.23.590724

**Authors:** Divyesh Joshi, Preeti Jindal, Ronak Shetty, Maneesha S. Inamdar

## Abstract

The Transforming Growth Factor β (TGFβ) signaling pathway is critical for survival, proliferation, and cell migration, and is tightly regulated during cardiovascular development. Smads, key effectors of TGFβ signaling, are sequestered by microtubules (MTs) and need to be released for pathway function. Independently, TGFβ signaling also stabilizes MTs. Molecular details and the *in vivo* relevance of this cross-regulation remain unclear, understanding which is important in complex biological processes such as cardiovascular development. Here, we use *rudhira/Breast Carcinoma Amplified Sequence 3 (BCAS3)*, a MT-associated, endothelium-restricted and developmentally essential proto-oncogene, as a pivot to decipher cellular mechanisms in bridging TGFβ signaling and MT stability. We show that Rudhira regulates TGFβ signaling *in vivo,* during mouse cardiovascular development, and in endothelial cells in culture. Rudhira associates with MTs and is essential for the activation and release of Smad2/3 from MTs. Consequently, Rudhira depletion attenuates Smad2/3- dependent TGFβ signaling thereby impairing cell migration. Interestingly, Rudhira is also a transcriptional target of Smad2/3-dependent TGFβ signaling essential for TGFβ-induced MT stability. Our study identifies an immediate early physical role and a slower, transcription-dependent role for Rudhira in cytoskeleton-TGFβ signaling crosstalk. These two phases of control could facilitate temporally-and spatially restricted targeting of the cytoskeleton and/or TGFβ signaling in vascular development and disease.

**Significance statement:** The developmental TGFβ pathway is essential for cell migration, cell-cell communication, adhesion, apoptosis, and matrix remodeling. Dysregulation of TGFβ signaling leads to aberrant vascular patterning and angiogenesis during mouse embryogenesis. Pathway activation involves phosphorylation and nuclear transport of Smads. Microtubules sequester Smads in the cytoplasm, thereby inhibiting TGFβ signaling. Conversely, TGFβ signaling stabilizes microtubules. However, the molecular components involved, and biological relevance of this cross-regulation remain unclear. We show that the oncoprotein Rudhira/BCAS3 facilitates Smad-MT dissociation upon ligand-mediated TGFβ receptor activation. Interestingly, Smad-dependent TGFβ signaling activation enhances rudhira transcription, essential for microtubule stabilization during cardiovascular development. This dual regulation of TGFβ signaling and microtubule stability by Rudhira allows sustained temporal control essential for development, and its dysregulation has pathological outcomes.

## Introduction

Cytoskeletal organization and growth factor signaling are mutually dependent and coordinate multiple cellular processes [1]. The multifunctional Transforming growth factor β (TGFβ) pathway regulates cell proliferation, differentiation, survival and migration and is sensitive to cytoskeletal rearrangements [1–3]. Depletion of TGFβ pathway components causes developmental abnormalities often leading to embryonic death [4]. Ligand binding activates the TGFβ receptor, causing receptor-mediated phosphorylation, followed by nuclear translocation of Smad proteins, resulting in transcriptional regulation of target gene expression [5]. Smads, the effectors of the TGFβ pathway, are competitively bound by several proteins and are a hub for regulation of TGFβ signaling and its intersecting pathways [6]. MTs bind to and sequester Smad2/3 and Smad4, thereby negatively regulating TGFβ signaling, which is relieved upon TGFβ stimulation [7]. Conversely, TGFβ signaling stabilizes MTs [8, 9] through an as yet unclear mechanism, suggestive of a feedback loop.

During development, TGFβ signaling is activated in endothelial cells (ECs) of the remodeling angiogenic endothelium and is critical for vascular development [10]. MT stability is also essential for angiogenic sprouting in ECs [11]. MTs can be stabilized in multiple ways within the cell, initially by RhoA-mDia actin bundling and thereafter by interaction with multi-protein complexes. Serum components, TGFβ and Lysophosphatidic acid (LPA), have been shown to stabilize MTs [8, 12]. While LPA stabilizes MTs within 30 min post-stimulation and generically [12], the effect of TGFβ signaling takes at least 2 hours [8] but is more restricted to and essential for endothelial cell function *in vivo*. Slow and restricted induction of MT stability by TGFβ as compared to the more generic and rapid function of LPA suggests a requirement for new transcription/translation which may be developmentally important. However, the molecular mechanisms and biological relevance of cytoskeletal interactions with TGFβ signaling remain elusive. Characterization of tissue-restricted regulators of the ubiquitous MT cytoskeleton may aid in defining its specialized context-dependent functions.

In this study, we address the role of the MT-interacting protein Rudhira in connecting TGFβ signaling and MT stability. Rudhira associates with MTs and intermediate filaments (IFs), stabilizes MTs and promotes directional cell migration [11, 13]. *Rudhira* knockout in mouse causes mid-gestation lethality due to aberrant cardiovascular patterning [14]. Transcriptome analysis showed that several processes were deregulated in the *rudhira* knockout, especially, a large number of TGFβ signaling pathway components [14]. This suggests that Rudhira may regulate TGFβ signaling for vascular development.

Here, we show that Rudhira-depleted cells have reduced Smad2/3 activation during mouse embryonic development, indicating that Rudhira is a positive regulator of TGFβ signaling. We demonstrate that Rudhira facilitates TGFβ-mediated release of Smad2/3 from MTs. Further, TGFβ signaling stabilizes MTs by inducing transcription, in a Rudhira-dependent manner. Thus, Rudhira bridges TGFβ signaling and MT stability, required for cardiovascular development. Ectopic and aberrant Rudhira expression is seen in several carcinomas and positively correlates with tumor metastatic potential [15–17]. Hence our study will aid understanding of TGFβ signaling and suggest novel ways to regulate it in development and disease.

## Results

### Rudhira depletion attenuates Smad2/3-dependent TGFβ signaling *in vivo* and *in vitro*

*Rudhira* is an essential gene, critical for mouse cardio-vascular development [14]. Ubiquitous (*rudh^-/-^*) or endothelial (*rudh^CKO^)* knockout of *rudhira* results in angiogenic defects and mid-gestation embryonic lethality [14]. Transcripts of several components of the TGFβ pathway are deregulated upon Rudhira depletion and the knockout phenocopies mutants of several TGFβ pathway components, whose primary phenotype is in the cardiovascular system [10]. Hence, we characterized the role of Rudhira in TGFβ signaling. Analysis of the molecular interaction, reaction and relation networks by KEGG pathway mapping indicated that multiple components of the TGFβ pathway were deregulated in the absence of Rudhira, affecting several processes important for cardiovascular development (Fig. 1A). To validate this, we used non-silencing (NS) control and rudhira knockdown (KD) saphenous vein endothelial cell lines (SVEC) as described before (Fig S1A, B) [14]. To test the pathway regulation at the level of receptors, we analysed transcript and basal as well as active protein levels. While we observed some reduction in transcript levels of *tgfbrI* and *tgfbrII* (Fig. S2A), the basal and active protein levels of TGFβRI remained unchanged in KD cells (Fig. S2B). Interestingly, a constitutively active form of TGFβRI (TGFβRIT204D), expressed under the CMV promoter, was also suppressed in Rudhira KD (overexpression achieved in NS ∼200 folds while in KD ∼80 folds) (Fig. S2C). TGFBR1 protein has a short half-life, while the mRNA is relatively stable [18]. Reduced amount of relatively stable RNA may be sufficient to produce adequate protein levels. These increased levels of active TGFβRI in KD were unable to restore Smad2 phosphorylation, unlike NS controls, which showed a clear increase (Fig. 1B). While important to note that Rudhira or its loss may regulate *tgfbr* transcript levels, these results support a receptor-independent role of Rudhira in regulating TGFβ pathway.

**Fig. 1.**
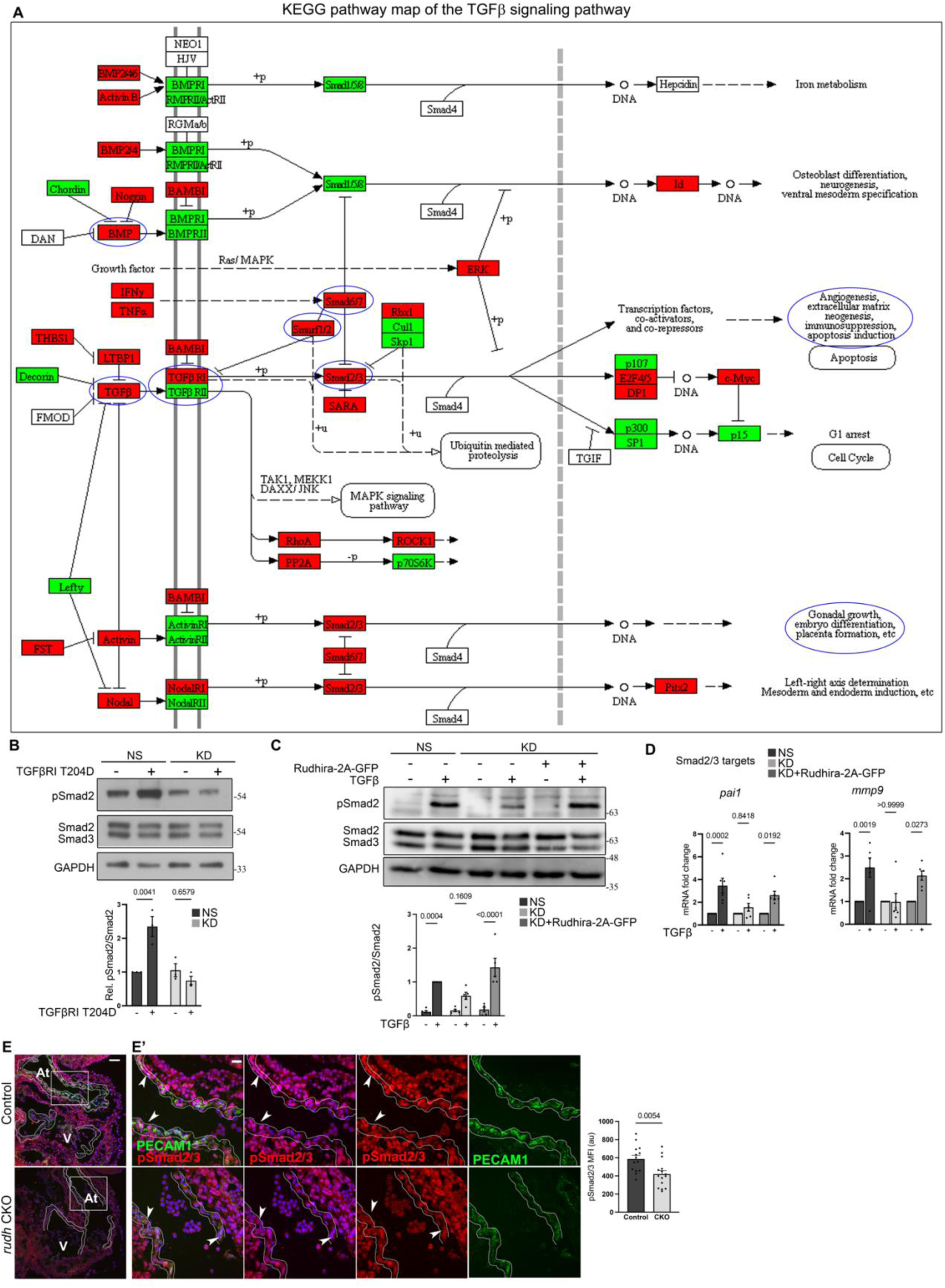
Rudhira depletion deregulates developmental endothelial TGFβ signaling. (A) KEGG pathway map indicating TGFβ pathway genes from KEGG database that are deregulated upon *rudhira* depletion in mouse embryonic yolk sac, based on [14]. Genes were mapped based on fold change in *rudhira*^-/-^ yolk sacs in comparison to control to understand pathway regulation. Green: downregulated, Red: upregulated. Blue circles indicate affected genes or processes known to be affected upon Rudhira depletion, from earlier studies. (B) NS and KD SVEC cells, transfected with constitutively active TGFβRI for 24h, were analyzed for Smad2 phosphorylation by Immunoblotting. Graph shows the quantitation of pSmad2/Smad2 levels (N= 3 independent experiments). (C, D) Control (NS), *rudhira* knockdown (KD) and rescued KD (KD + Rudhira-2A-GFP) cells were kept untreated or treated with TGFβ and used for various assays, as indicated. (C) Immunoblot for Smad2/3 and pSmad2. Graph shows the quantitation of pSmad2/Smad2 levels (N= 3 independent experiments). (D) qRT-PCR analysis of *mmp9* and *pai1*, known Smad2/3 targets in ECs (N= 3 independent experiments). (E, E’) Immunostaining for phosphorylated Smad2/3 in control and *rudhira* conditional knockout *(rudh^CKO^)* embryos at E10.5. Boxed region in the embryonic heart in (E) marking the endocardium and the myocardium are magnified in (E’) (N= 3 embryos for each genotype). White dotted line indicates the endocardium. Graph shows the quantitation of pSmad2/3 levels (N= 3 independent experiments). Statistical analysis was performed using two-way ANOVA.(A, B, C, D) and student’s t-test (E). Error bars indicate standard error of mean (SEM). *p<0.05, **p<0.01, ***p<0.001. Scale bar: (E) 100 µm; (E’) 20 μm.

To probe further into the molecular mechanism by which Rudhira controls TGFβ signaling, we used *rudhira* knockout mice and the KD endothelial cell line. Upon TGFβ-stimulation ECs in culture showed reduced Smad2 activation in the KD as compared to NS control (Fig. 1C). Further, qRT-PCR analysis showed that transcript levels of Smad2/3 targets, namely *mmp9* and *pai1*, did not increase significantly in KD, whereas NS control showed >2-fold increase for both (Fig. 1D). Expectedly, partial restoration of Rudhira level in KD cells restored Smad2 activation and target gene expression (Fig. 1C, D). The canonical TGFβ pathway also responds to BMP4 activation. However, upon stimulation with BMP4 there was no significant difference in the levels of pSmad1/5/8 between NS and KD (Fig. S2D). These data indicate that Rudhira is required for TGFβ-dependent Smad2 phosphorylation and pathway activation.

Endothelial depletion of Rudhira (*rudh^CKO^)* is sufficient to cause vascular defects [14] and functional TGFβ signaling is essential for vascular development [19]. Expectedly, *rudh^CKO^* embryos showed a concomitant loss of pSmad2/3 in the embryonic endocardium (Fig. 1E, E’), without affecting the total Smad2/3 level (Fig. S2E, E’). Thus, Rudhira regulates TGFβ signaling in a Smad2/3-dependent manner *in vivo*, which is essential for angiogenesis and endothelial cell migration during cardio-vascular development.

### Rudhira is required for TGFβ mediated cell migration

As Rudhira is essential for cell migration, we tested the effect of altered TGFβ pathway activity in Rudhira KD endothelial cells (SVEC) in a transwell migration assay. We also validated our results in HEK293 cells as these allow a high efficiency of transfection and Rudhira overexpression, for migration assays. TGFβ addition did not enhance migration in *rudhira* knockdown (KD) cells as compared to controls (NS and rescued KD) (Fig. 2A). In addition, ectopic Rudhira expression in HEK293 cells caused increase in migration rate as compared to the control, consistent with our earlier study [13] (Fig. 2B,C).

**Fig. 2.**
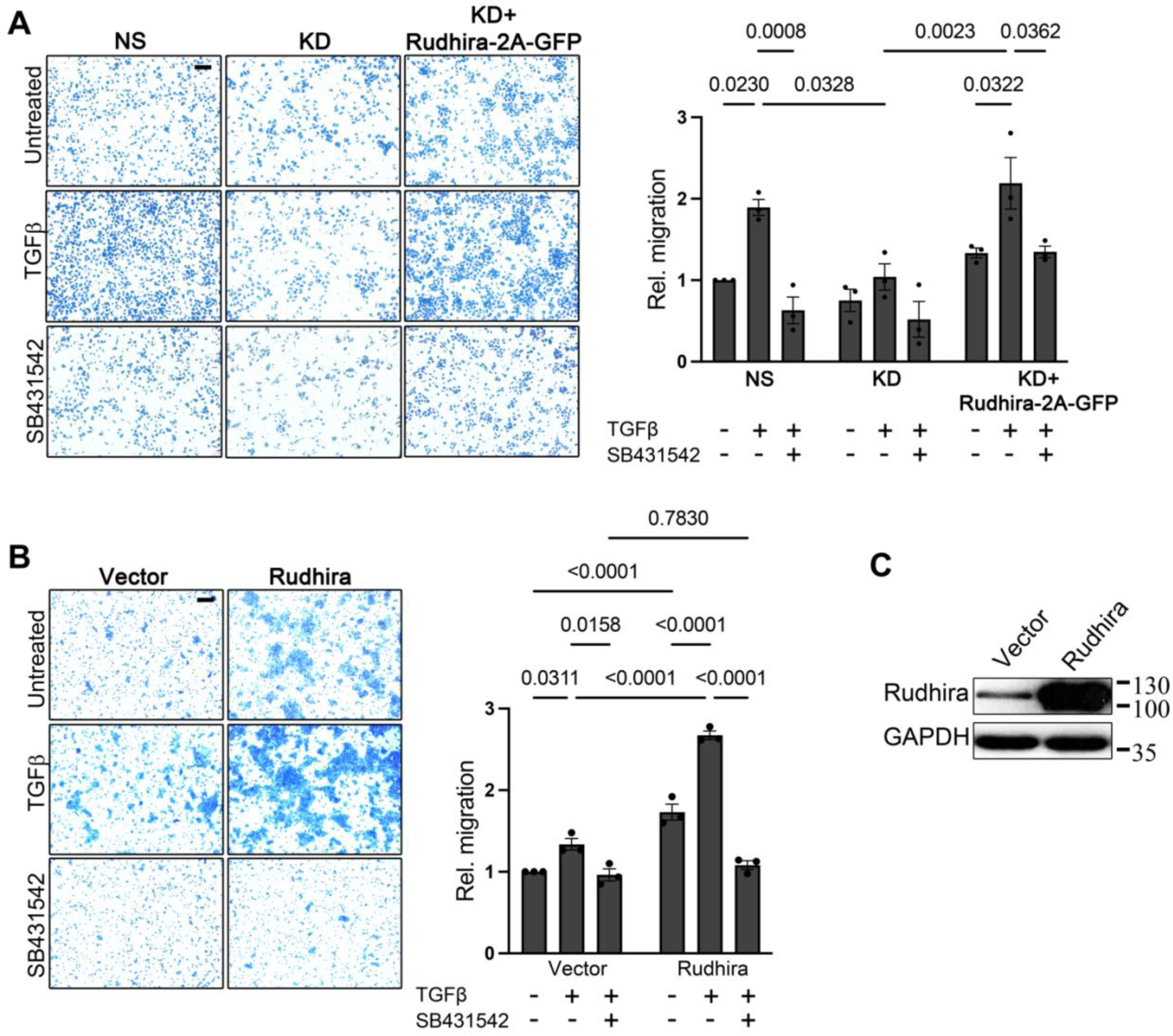
Rudhira functions downstream of TGFβ receptor activation to promote TGFβ-dependent cell migration. (A) Control (NS), *rudhira* knockdown (KD) and rescued KD (KD + Rudhira-2A-GFP) cells were analyzed for migration rates with or without TGFβ using a transwell migration assay. Graph shows the extent of cell migration as measured by crystal violet absorbance (N= 3 independent experiments). (B, C) HEK293 cells transfected with vector alone or Rudhira were analyzed for migration rates using a transwell migration assay in various conditions, as indicated. Graph shows the extent of cell migration as measured by crystal violet absorbance (N= 3 independent experiments). Statistical analysis was performed using one-way ANOVA. Error bars indicate SEM. Scale bar: (A, B) 100 μm. *p<0.05, **p<0.01, ***p<0.001.

TGFβ supplementation of Rudhira overexpressing cells dramatically enhanced the rate of cell migration (∼2.6 fold), suggesting a direct correlation between Rudhira levels and pathway function (Fig. 1C, D). Serum contains TGFβ in a concentration range of 10-40 ng/ml [20] and we used 5% FBS with or without additional TGFβ in the transwell migration assay, since purified TGFβ alone is unable to induce migration. The ratio of increase in migration rates with or without exogenous TGFβ addition was similar in both the control and Rudhira-overexpressing cells (∼1.5 fold). Interestingly, TGFβ pathway-inhibition by SB431542 (SB) restored the effect of Rudhira overexpression (enhancement in cell migration rate) to control levels (Fig. 2B, S3A) suggesting that Rudhira function is regulated by the TGFβ pathway downstream of receptor activation. In addition to the multiple molecular pathways that function in concert to control cell migration [21], our data support a role for Rudhira in TGFβ-dependent cell migration.

### Rudhira regulates TGFβ signaling by promoting the release of Smad2/3 from MTs

Since Rudhira localizes to the cytoskeletal components, namely microtubules and intermediate filaments [13], we hypothesized that it regulates TGFβ signaling at the cytoskeleton. However, we could not detect any interaction between Rudhira and Smad2/3 under the conditions tested (Fig. 3A), suggesting that the effect on the pathway may be indirect, possibly through MTs. TGFβ signaling is controlled at multiple levels, including cytoskeletal regulation at MTs, which interact with and sequester Smad2/3 thereby inhibiting the TGFβ pathway [7]. TGFβ stimulation releases Smads from MT sequestration in multiple cell types including endothelial cells, facilitating its activation and nuclear translocation [7]. Expectedly, TGFβ stimulation in ECs (confirmed by increase in pSmad2 levels) reduced Smad-MT binding as seen by co-immunoprecipitation for β-Tubulin and immunoblotting for Smad2 (Fig. 3B).

**Fig. 3.**
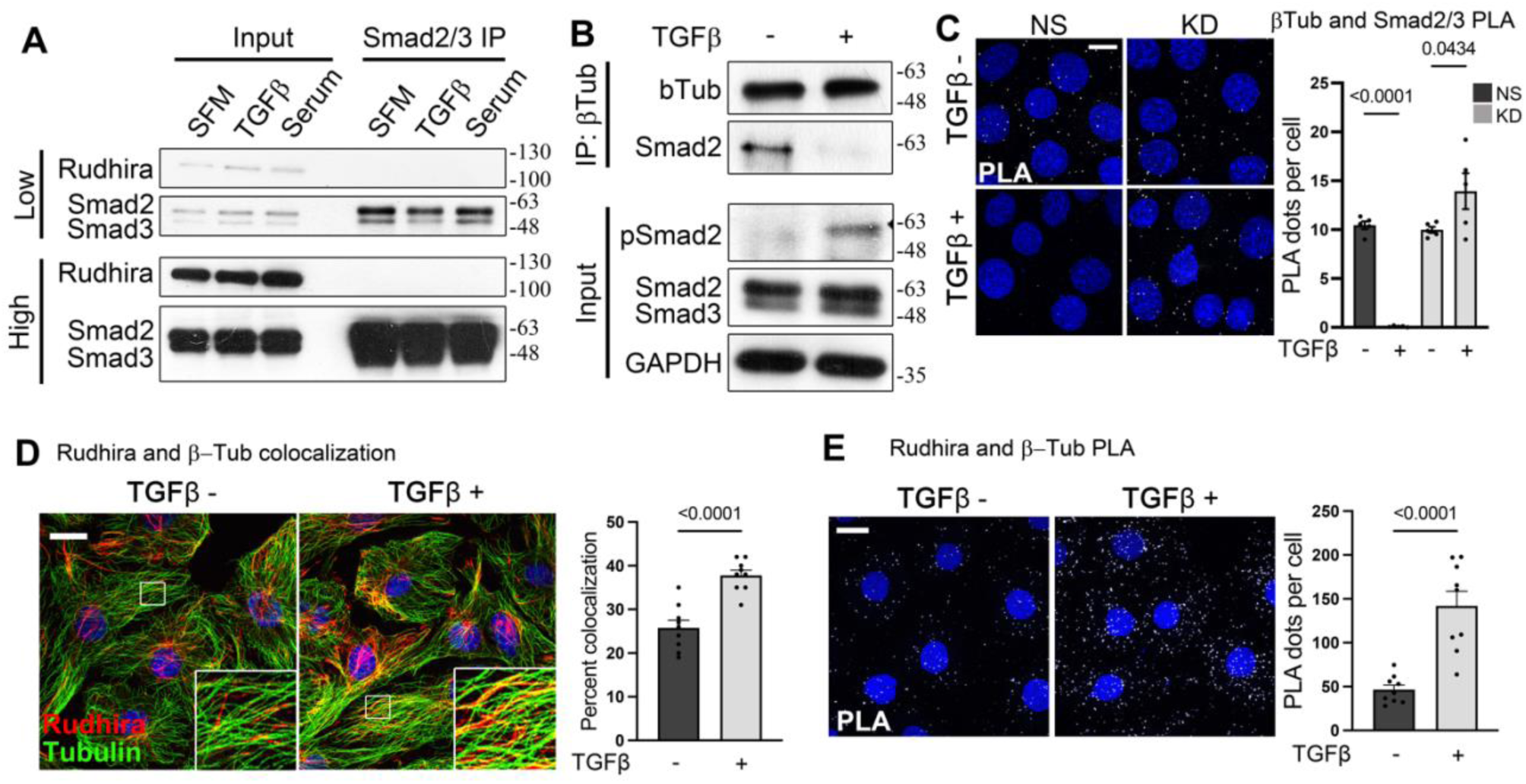
Rudhira is essential for the release of Smads from microtubules for TGFβ pathway activation. (A) Analysis of interaction between Rudhira, Smad2/3 and MTs by co-immunoprecipitation and immunoblotting, as indicated (N= 3 independent experiments). (B) Loss of Smad-MT interaction on TGFβ stimulation in SVEC cells (N= 3 independent experiments). (C) NS and KD cells were analyzed for MT and Smad2/3 interaction by PLA for Tubulin and Smad2/3 with or without TGFβ treatment. PLA dots represent Smad2/3 and β-tubulin interaction. Graph shows the quantitation of PLA dots per cell (quantitation from >80 cells for two knockdown cell lines from N= 3 independent experiments; also see Fig. S3). (D, E) Association of Rudhira and MTs was analyzed by immunostaining (D) or Proximity Ligation Assay (PLA) (E) for Rudhira and Tubulin with or without TGFβ stimulation in SVEC. PLA dots represent Rudhira and β-tubulin interaction. Graph (D) shows the quantitation of colocalization percentage (quantitation from ∼40 cells from 9 images each with or without TGFβ respectively from N= 3 independent experiments) and graph (E) shows the quantitation of PLA dots per cell (quantitation from 86 or 68 cells from 9 images each with or without TGFβ respectively from N= 3 independent experiments). Statistical analysis was performed using one-way ANOVA. Error bars indicate SEM. Scale bar: (C) 20 μm; (D, E) 10 µm. *p<0.05, **p<0.01, ***p<0.001.

To assess whether Rudhira is required for TGFβ-dependent release of Smad from MTs, we tested for β-Tubulin and Smad interaction upon TGFβ stimulation in NS and KD cells by performing *in situ* Proximity Ligation assay (PLA) [22]. While NS cells showed reduced Smad-MT interaction, KD cells retained Smad-MT interaction even upon TGFβ stimulation in multiple KD cell lines (Fig. 3C, Fig. S3B). Interestingly, TGFβ stimulation resulted in an increase in Rudhira-MT association in wild type cells, detected by colocalization (Fig. 3D) and confirmed by PLA (Fig. 3E). Thus TGFβ-stimulated Rudhira-MT association makes MTs unavailable for Smad binding. However, MTs are abundant and present as subsets, differing in their Tubulin isotype and isoform composition, post translational modifications (PTMs) and stability [23], which may also govern their interactions with other molecules. It is unlikely that Rudhira binds to all MTs present. Also, it is likely that only a subset of MTs may bind and sequester Smads.

### *Rudhira* is a transcriptional target of TGFβ signaling and is essential for MT stability

The TGFβ pathway is auto-regulatory, due to which several pathway components are induced or inhibited [24, 25]. The *rudhira* promoter harbors putative Smad-binding elements (SBEs) (Fig. S4A). It also harbors putative binding sites of major TGFβ pathway regulated transcription factors (TFBS) (Fig. S4B, Table S2). Expectedly, TGFβ stimulation increased *rudhira* mRNA and protein levels (Fig. 4A, B, C). Knockdown of *smad2* or *smad3* resulted in significant reduction of Rudhira levels, indicating that *rudhira* is a Smad2/3-dependent TGFβ target (Fig. 4D). Blocking transcription with Actinomycin D (AcD) or translation using Cycloheximide (CHX) prevented increase in *rudhira* levels upon TGFβ stimulation, confirming that *rudhira* is a transcriptional target of the Smad2/3-dependent TGFβ pathway (Fig. 5A, S4C). However, additional investigation is required to confirm whether *rudhira* is a direct target of Smad2/3-dependent TGFβ pathway.

**Fig. 4.**
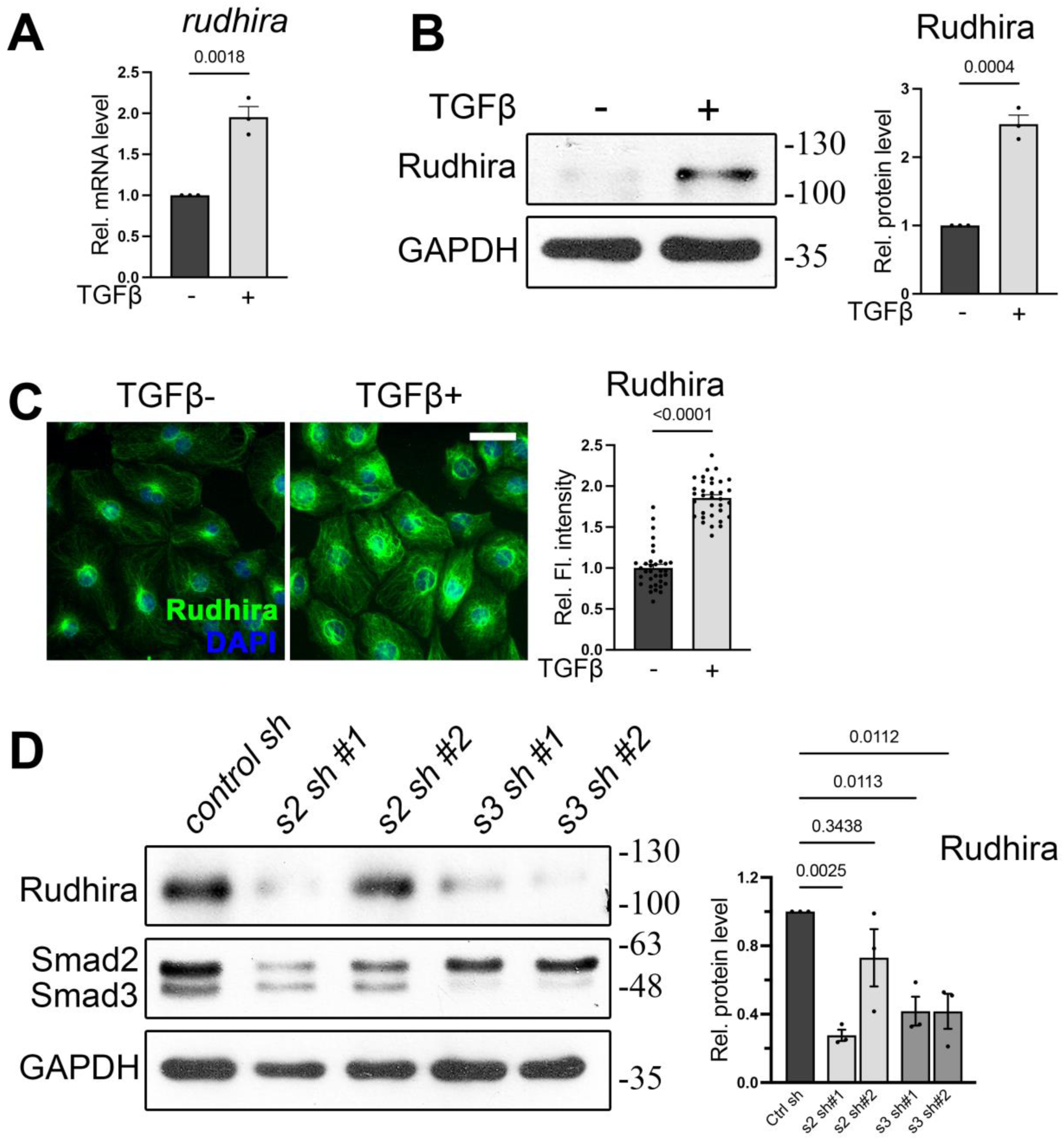
Rudhira is a Smad2/3-dependent target of TGFβ signaling. (A-C) TGFβ stimulation followed by qRT-PCR (A), immunoblot (B) or immunostaining (C) analysis (quantitation from 34 cells in each condition) of Rudhira levels in SVEC cells. Graphs in (B) and (C) show the quantitation of Rudhira levels with or without TGFβ (N= 3 independent experiments). (D) Analysis of Rudhira levels upon *smad2* or *smad3* knockdown in HEK293T cells by immunoblotting (N= 3 independent experiments). Statistical analysis was performed using one-way ANOVA. Error bars indicate SEM. Scale bar: (C) 10 µm. *p<0.05, **p<0.01, ***p<0.001.

**Fig. 5.**
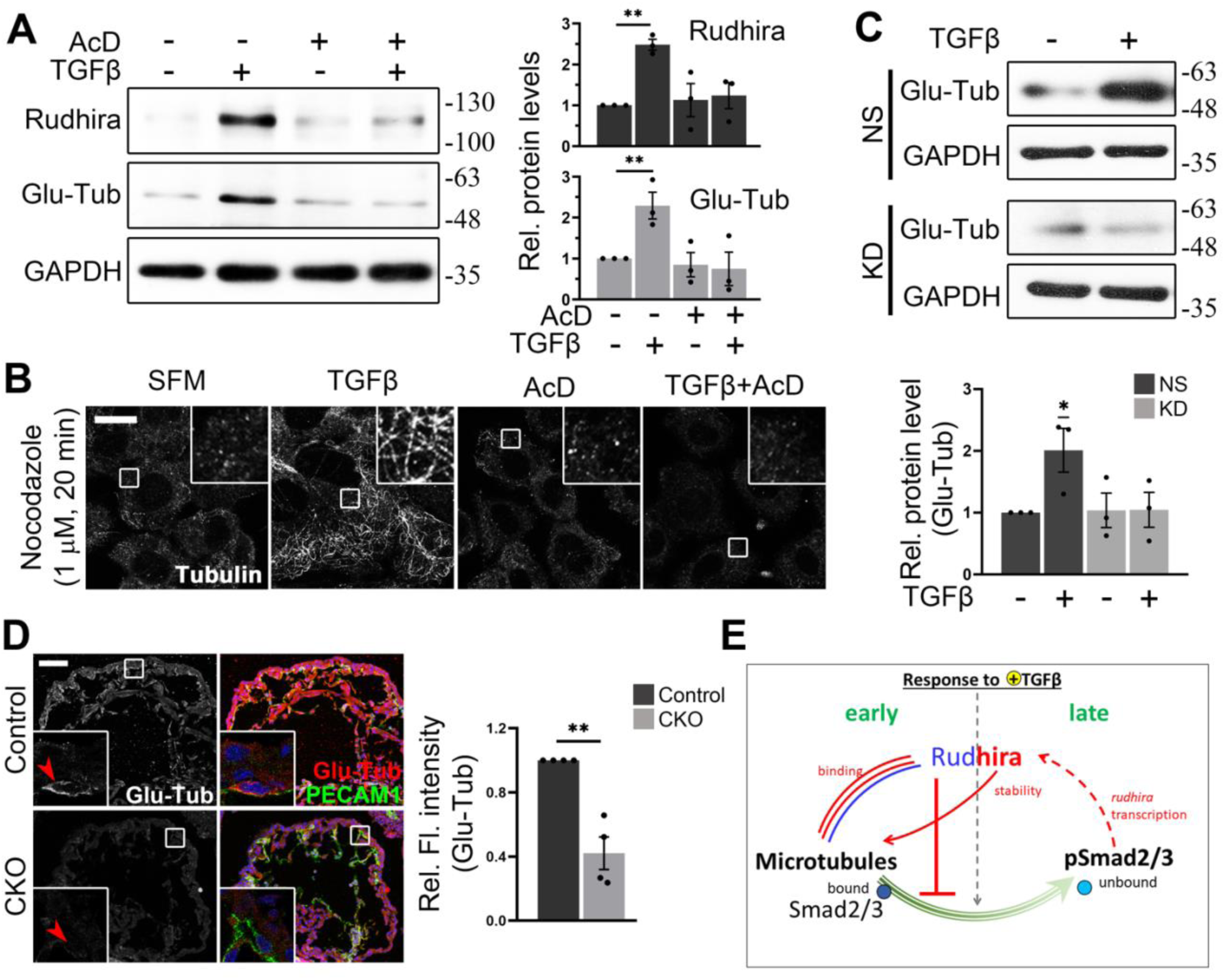
TGFβ-dependent *rudhira* transcription stabilizes MTs. (A) Analysis of Rudhira levels and MT stability (marked by Glu-Tubulin) upon treatments as indicated, by immunoblotting. Graphs show the quantitation of Rudhira or Glu-Tub levels (N= 3 independent experiments). (B) Analysis of MT resistance to Nocodazole mediated depolymerization at indicated dosage and time (N= 30 cells). (C) MT stability in serum starved NS or KD SVEC cells kept untreated or treated with TGFβ was analyzed by immunoblot for Glu-Tubulin. Graph shows the quantitation of Glu-Tub levels (N= 3 independent experiments). (D) Immunostaining for Glu-Tubulin in control and *rudhira* conditional knockout *(rudh^CKO^)* embryos at E10.5. Endocardium in the heart is marked by PECAM1. Boxed region in the embryonic heart is magnified in the insets. Red arrowheads in the insets mark the PECAM1 positive cells (N= 4 embryos for each genotype). Graph shows the quantitation of Glu-Tub fluorescence intensity in the endocardium. (E) Model depicting cellular and molecular response to TGFaddition. Rudhira has an early and a late effect on MTs. Previously known information about Rudhira is shown in blue. Events identified in this study are shown in red or pink. Parallel lines indicate protein-protein interaction. Solid lines indicate direct effects. Dotted lines indicate effects that may be direct or indirect. Arrows indicate positive regulation. Bar-headed line indicates inhibition. Statistical analysis was performed using one-way ANOVA. Error bars indicate SEM. Scale bar: (B) 10 µm; (D) 100 µm. *p<0.05, **p<0.01, ***p<0.001.

Rudhira and TGFβ signaling, both can induce MT stabilization [8, 11]. Although the mechanism is unclear, PTMs and/or transcription may control TGFβ-induced MT stability [8]. Since *in vivo* studies indicated that *rudhira* deletion severely affects the TGFβ pathway [14], we tested whether TGFβ-induced transcription is required for MT stability. TGFβ stimulation results in a slow enhancement of MT stability over a period of several hours, allowing sufficient time for transcriptional regulation [8]. Interestingly, we find that blocking transcription or translation, prevented TGFβ-dependent MT stabilization, as detected by Glu-tubulin levels (Fig. 5A, S4C), correlating well with Rudhira levels and validated by increased sensitivity to Nocodazole-mediated MT depolymerization (Fig. 5B). These data show that TGFβ induced MT stability is transcription-dependent.

### TGFβ signaling stabilizes MTs in a Rudhira-dependent manner

Since *rudhira* transcription is dependent on TGFβ signaling and Rudhira is required for MT stability, we further tested whether TGFβ-induced MT stability is Rudhira-dependent. Serum starvation resulted in low detectable levels of Glu-tubulin in NS and KD cells (Fig. 5C). Stimulation with TGFβ or serum led to increased Glu-tubulin level in NS but not in KD cells showing that TGFβ-induced MT stability is dependent on Rudhira (Fig. 5C, D). To test whether Rudhira stabilizes MTs *in vivo* also, we analyzed the effect of loss of Rudhira on MT stability in the developing endocardium. Interestingly, along with reduced pSmads levels observed earlier (Fig. 1E, E’), we found that Glu-tubulin positive stable MTs were fewer in *rudh^CKO^* embryonic endocardium (Fig. 5D), supporting a role of Rudhira in regulation of TGFβ signaling and MT stability during cardiovascular development. Combined, our study suggests that Rudhira primarily affects MTs to modulate their Smad-binding property and downstream TGFβ signaling. Consequently, Smad2/3-dependent TGFβ signaling induces *rudhira* transcription leading to MT stabilization essential for cardiovascular remodeling during development (Fig. 5E).

## Discussion

Growth factor signaling and cytoskeletal remodeling are mutually dependent processes that require complex context-dependent molecular crosstalk [1]. Specifically, the developmentally essential, tissue-restricted TGFβ signaling pathway, regulates and is regulated by the MT cytoskeleton. However, the underlying molecular mechanism and *in vivo* relevance of this cross-regulation remain elusive. In this study, we probed the role of Rudhira in the cytoskeletal control of TGFβ signaling and, in turn, its effect on MT properties. Our study identifies a dual role of Rudhira during vascular development-a n e a r l y r o l e in MT-mediated regulation of Smad2/3 and a late role as a transcriptional target of TGFβ signaling required for MT stability.

While several processes important for cardiovascular development were deregulated in *rudhira^-/-^* embryos, the TGFβ pathway was primarily affected [14]. TGFβ signaling is a complex pathway and is regulated at multiple levels by various feedback loops. One of the complicated yet delicate feedback circuits is the regulation of receptor levels and activity [25]. Rudhira depletion reduces transcript level without altering basal and active protein level of TGFBRI. This disparity in the transcript and protein level may arise from the fact that while the *tgfbr* transcript is very stable, the protein is quite unstable, and lower transcript levels may still be sufficient for generating adequate protein levels. Such a phenomenon is not unusual, particularly for membrane proteins, where mRNA levels may not correlate with protein levels. Rudhira, in a manner not yet understood, may also be a part of a broader feedback pathway as it further negatively regulates levels of TGFβ pathway inhibitors such as Smurfs and Smad7 [14], suggesting that additional regulatory mechanisms may operate.

TGFβ signaling and Rudhira promote cell migration in a synergistic and inter-dependent manner. Mechanistically, Rudhira promotes TGFβ-dependent release of Smads from MTs likely due to increased Rudhira-MT association. This could allow pathway activation and result in enhanced cell migration. Additional, confounding factors downstream of the autoregulatory TGFβ pathway or pathway targets, as described above, may also contribute to Rudhira dependent cell migration in response to TGFβ. Converse regulation of Rudhira-MT and Smad-MT interaction is indicative of probable competition between Smads and Rudhira for MT binding, which merits further investigation. Binding of Rudhira to MTs may also alter them in a way that makes MTs unavailable for Smad binding. Since TGFβ stimulation is known to stabilize microtubules, we hypothesize that TGFβ stimulation increases Rudhira binding to stable microtubules. While unclear molecular events involved in release of Smads from MTs, and MT stabilization by Rudhira make it difficult to assign causal roles, this study supports the hypothesis that dissociation of Smads from MTs is essential for Smad activation. In addition, though MTs are ubiquitously and abundantly present, our data suggest that only a few MTs participate in Smad sequestration. This is in concordance to our earlier study which shows that Rudhira associates preferentially with stable MTs [11].

The TGFβ signaling pathway is autoregulatory and many of its components are also targets of the pathway [24]. The *rudhira* promoter harbors Smad Binding Elements (SBEs). We show that Rudhira, in addition to being a regulator of the TGFβ pathway, is also a transcriptional target. However, further analysis is required to identify the specific mechanisms by which Smad-dependent, TGFβ-induced, *rudhira* transcription is regulated during development. In contrast to this positive feedback, TGFβ pathway is also negatively self-regulated, allowing for return to basal state amenable for reactivation. This self-suppression could possibly involve increased turnover of receptors and Smad2/3, and increased expression of inhibitory Smads, that may recover responsiveness to TGFβ stimulation. Additionally, in the context of MT-dependent Smad2/3 inhibition, the still short turnover time of stable MTs (several minutes to hours) may also promote quick return to resting state. These interesting possibilities can now be tested.

*Rudhira* expression is tightly regulated and restricted to the remodeling endothelium during development [14]. Mechanistically, Rudhira promotes MT stability for angiogenic remodeling [11]. Independent studies show that TGFβ signaling, and MT reorganization are essential for cardiovascular development [26–28]. Deletion of *smad2/3* or *tubulins* leads to embryonic lethality with severely defective embryonic and extra-embryonic vasculature [29]. However, the role of MT stability for angiogenic sprouting *in vivo* and the underlying molecular mechanisms remain elusive. Our study suggests that endothelial cytoskeleton remodeling by Rudhira and activation of TGFβ signaling during cardiovascular development are inter-dependent and cross-regulated. TGFβ signaling may lead to restricted induction of MT stability by inducing the expression of MT-stabilizing developmentally regulated proteins like Rudhira. Additional cytoskeletal components such as Vimentin that interacts with Rudhira, and is also a target of TGFβ signaling, are likely to be involved [11, 13]. Our analysis will help gain better insight into cellular processes such as migration, which require maintenance of cytoskeletal stability and active signaling for extended periods. An added advantage will be the ability to decipher deregulated migration such as in tumor metastasis. While targeting MT-Smad interaction would allow regulation of the TGFβ pathway, the ubiquitous nature of these molecules is likely to result in widespread and possibly undesirable effects of therapeutic intervention. Rudhira expression, being more restricted, could provide a suitable target to regulate developmental and pathological TGFβ signaling.

## Materials and methods

### Animal experiments

Animal experiments were performed as mentioned earlier [14]. All animal experimental protocols were approved by the Institutional Animal Ethics Committee (IAEC) of JNCASR (Project number MSI006. All animals were maintained, and experiments were performed according to the guidelines of the animal ethics committee of JNCASR. *Rudhira* floxed mice and conditional knockout mice (*Tek-Cre* mediated endothelial deletion, *rudhira^flox/flox^; TekCre^+^*, abbreviated to *rudh^CKO^*) were validated by genotyping.

### Cell culture, transfections, and small molecule treatments

Cell line sources and generation is reported elsewhere [14]. In brief, *rudhira* shRNA vectors (715, 716), and scrambled (non-silencing) control vector (TR30015) (Origene, USA) were microporated into SVEC (mouse endothelial cell line) and selected for stable line generation. Control cell line is referred to as ‘NS’ and *rudhira* knockdown as ‘KD’. For Rudhira overexpression and rescue line, the plasmid constructs used were GFP-2A-GFP and Rudh-2A-GFP which are described elsewhere [13]. Constitutively active TGFβRI (pcDNA3-ALK5 T204D, 80877) was purchased from addgene for transfection of SVEC NS and KD cells. Control and shRNA vectors for s*mad2* or *smad3* knockdown) were purchased from ShRNA Resource Centre (Department of Microbiology and Cell Biology, Indian Institute of Science, Bangalore, India) and were from the SIGMA Library of shRNAs for Human gene products. The shRNAs were transfected in HEK293T cells and selected in 1 µg/ml Puromycin for 7 days. Details of the oligonucleotide sequence of the shRNAs are provided in Supplementary Table S3. HEK293, HEK293T cells were transfected using Calcium Phosphate method while SVEC cells were transfected using Lipofectamine 2000 (ThermoFisher Scientific, USA). All small molecules were from Sigma Chemical Co., USA, and treatments were performed in DMEM, unless otherwise indicated. Cells were washed three times in PBS and serum starved for 12h. Thereafter, cells were kept untreated or treated with 10 ng/ml of TGFβ (Fig. 1C) or BMP4 (Fig. S2D) for 2 h and then taken for immunoblotting. For gene expression analysis, cells were induced with 10 ng/ml of TGFβ for 24 to 48 h (Figs. 1D, 4A, B, C, S3C). SB431542 was used at a concentration of 10 µM, as indicated (Fig. 2A, B and S3A). For Transwell migration assays, small molecule dilutions were prepared in 5% FBS containing DMEM (Fig. 2A, B). For interaction studies cells were serum starved for 12h and thereafter kept untreated or treated with 10 ng/ml of TGFβ or 10% FBS for 2 h and then taken for PLA, immunostaining or western blotting, as desired (Fig. 3, S3A, B). For TGFβ-dependent *rudhira* transcription and MT stability assays, cells were serum starved for 48h in serum-free medium (SFM) containing 5 mg/ml BSA and 20mM HEPES (pH 7.4) in DMEM. Thereafter, various treatments were performed for 7h (Fig. 5A, B, C, S4C, D). TGFβ (10 ng/ml), Actinomycin D (AcD, 10 µM), Cycloheximide (50 µg/ml) were diluted in SFM.

### Quantitative RT-PCR (qRT-PCR)

RNA was isolated using TRIzol reagent (Invitrogen). Reverse transcription was performed using 2 μg of DNase treated RNA and Superscript II (Invitrogen, Carlsbad, CA) according to manufacturer’s instructions. Quantitative RT-PCR (qRT-PCR) was carried out using EvaGreen (BIO-RAD, CA) in Biorad-CFX96 Thermal cycler (BIO-RAD, CA). Details of primers used are provided in Supplementary Table S1.

### Immunostaining, Immunohistochemistry, Fluorescence Microscopy and Analysis

Yolk sac or embryos dissected at E10.5 were fixed in 4% paraformaldehyde and processed for Cryosectioning (embryos) and immunostaining using standard procedures. Control and knockout embryos were identified by embryonic tail genotyping. Cells were fixed in 4% paraformaldehyde at room temperature or 100% methanol at-20 ℃ and processed for immunostaining using standard procedures [30]. Primary antibodies used were against PECAM1 (CD31) (BD Biosciences, USA), Rudhira, Smad2/3, pSmad2, pSmad2/3 (Cell Signaling Technology), β-Tubulin (DSHB, Iowa), Glu-Tubulin (Abcam), GFP (Invitrogen). Secondary antibodies were conjugated with Alexa-Fluor 488 or Alexa-Fluor 568 or Alexa-Fluor 633 (Molecular Probes). Bright field and phase contrast microscopy were performed using an inverted (IX70, Olympus) microscope. Confocal microscope, LSM 880 with airy scan, Zeiss, was used for fluorescence microscopy. All images in a set were adjusted equally for brightness and contrast using Adobe Photoshop CS2, where required.

### Immunoblot analysis and co-immunoprecipitation

50 µg whole cell lysate was used for western blot analysis by standard protocols. Blots were cut into strips and incubated with primary antibodies as indicated: Smad2/3, pSmad2, Smad1/5/8, pSmad1/5/8 (Cell Signaling Technology), GAPDH (Sigma Chemical Co., USA), Glu-Tubulin, Ubiquitin, TGFβRI, pTGFβRI (Abcam, USA), β-Tubulin (DSHB, Iowa), Rudhira (Bethyl Labs, USA). HRP conjugated secondary antibodies against appropriate species were used and signal developed by using Clarity Western ECL substrate (Biorad, USA). For co-immunoprecipitation, 500 µg whole cell lysate and 2 µg of the desired antibody were used. Complexes were captured on Protein G-Agarose beads and analyzed by immunoblotting.

### Transwell migration assay

The assays and quantitation were carried out as mentioned earlier [13]. Briefly, 24 h after transfection with desired plasmid vectors, cells were serum-starved for 12 h and 20000 HEK cells or 40000 SVEC were plated onto the upper chamber of the transwell filter inserts with 8 μm pore size, 24-well format (Costar, USA). 10% FBS containing medium (with desired small molecules) was added to the lower chamber to serve as a chemo-attractant. After 24 h, cells were fixed in 4% paraformaldehyde for 10 min at room temperature. Cells on the top of the filter were removed using a cotton swab. Cells that had migrated to the bottom were fixed and stained with 0.5% crystal violet for 10 min at room temperature. The dye was extractedin methanol and absorbance measured spectrophotometrically at 570 nm.

### *In situ* Proximity Ligation Assay (PLA or Duolink assay)

*In situ* PLA (Proximity Ligation Assay) reaction was performed on SVEC cell lines. The cells were cultured, fixed, permeabilised and stained with primary antibodies for Smad2/3, β-Tubulin or Rudhira, as desired. Thereafter, the protocol for PLA as recommended by manufacturer (Duolink, USA) was followed. Post PLA, nuclei were counterstained with DAPI.

### Rudhira promoter *in silico* analysis

Smad-binding elements or motifs (SBEs) and other transcription factor binding sites (TFBS) were identified from experimental studies, available in JASPAR database (https://jaspar.genereg.net/). PSCAN software (http://159.149.160.88/pscan) was used for motif scanning in mouse and human *rudhira/BCAS3* gene promoters (1 kb sequence upstream of the transcription start site) using matrices obtained from JASPAR bioinformatics tool, as indicated. (Supplementary Table S2).

### Quantification and Statistical analyses

Statistical analyses were performed using One Way ANOVA in the Data Analysis package in Microsoft Excel, and two-way ANOVA and one-sample t-test in GraphPad Prism. p<0.05 was considered significant.

## Acknowledgements

We thank Aksah Sam for maintaining mouse stocks; Developmental Studies Hybridoma Bank, University of Iowa, USA for some antibodies; JNCASR Imaging facility, JNCASR Animal Facility, NCBS Animal facility for access and Inamdar laboratory members for fruitful discussions. We are thankful to Arghakusum Das for repeating the experiment for Figure 4A. This work was funded by grants from the Department of Biotechnology, Government of India (Sanction no. BT/PR11246/BRB/10/644/2008 dated 29.09.2009), JC Bose fellowship (Grant no. JCB/2019/000020), Department of Science and Technology, Government of India, the Wellcome Trust, UK (094879/B/10/Z) and intramural funds from Jawaharlal Nehru Centre for Advanced Scientific Research, India.

## Author Contributions

MSI conceived of the project and directed the work. DJ, PJ, RS, MSI performed and analyzed all the experiments. DJ, PJ, MSI wrote the manuscript. Authors have no competing interests.

## Supplementary Figures and Figure legends

**Fig. S1.**
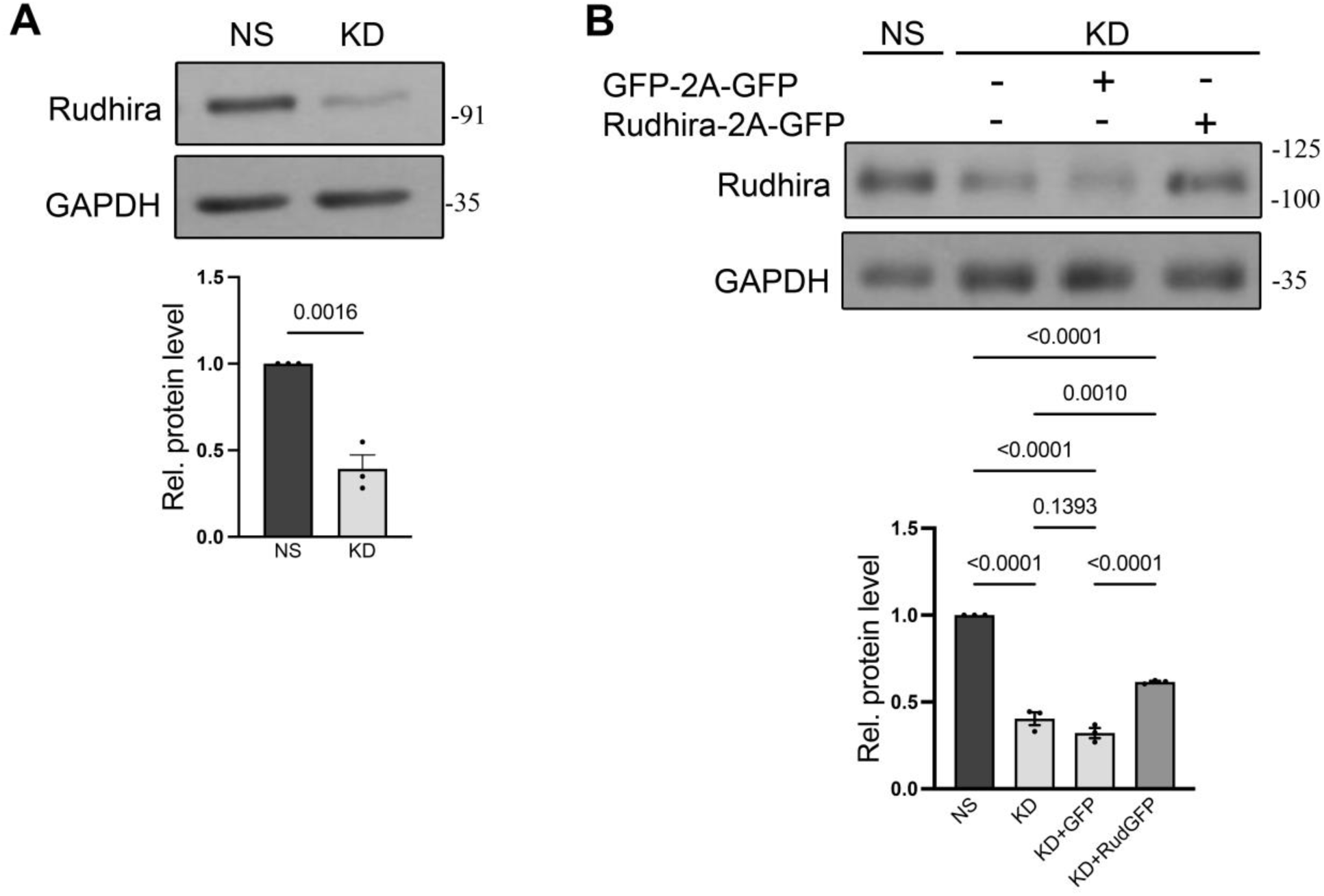
(related to Fig. 1). Cell line validation. (A, B) Validation of Rudhira knockdown and restoration in SVEC cells by immunoblotting. Statistical analysis was performed using one-sample t-test. Error bars indicate standard error of mean (SEM). *p<0.05, **p<0.01,***p<0.001.

**Fig. S2.**
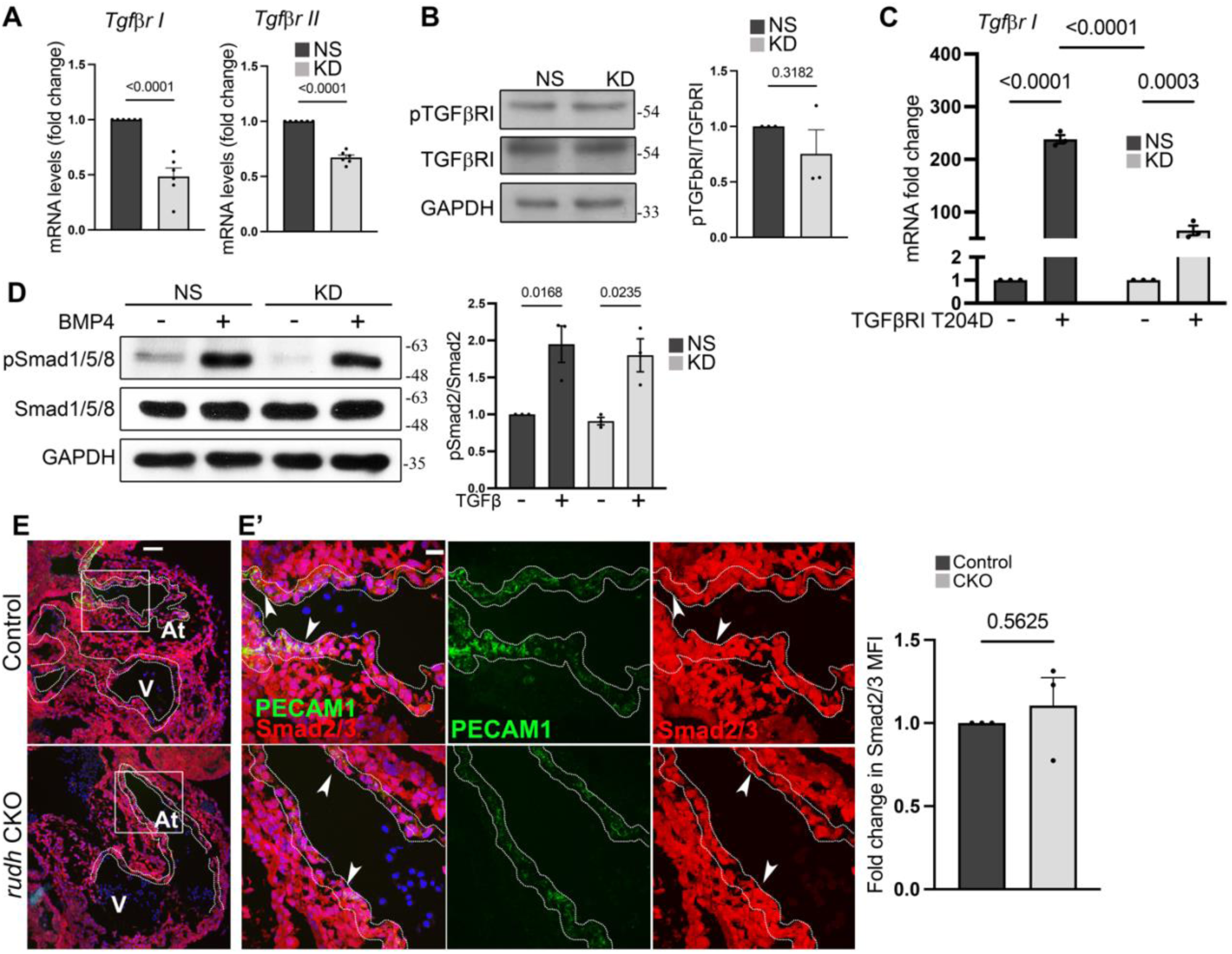
(related to Fig. 1). Rudhira acts downstream to TGFβ receptors. (A) qRT-PCR analysis of *tgfβrI* and *tgfβrII* in Rudhira knockdown endothelial cells (N= 3 independent experiments). (B) TGFβRI phosphorylation was analyzed using immunoblot technique in SVEC NS and KD. Graph shows the quantitation of pTGFβRI/TGFβRI levels (N= 3 independent experiments). (C) NS and KD SVEC cells, transfected with constitutively active TGFβRI for 24h, were analyzed for total TGFβRI level by qRT-PCR. (D) SVEC control (NS) and knockdown (KD) cells were stimulated by BMP4 and used for immunoblotting of pSmad1/5/8. Graph shows the quantitation of pSmad1/5/8 by Smad1/5/8 levels (N= 3 independent experiments). (E, E’) Immunostaining for total Smad2/3 in control and rudhira conditional knockout (rudhCKO) embryos at E10.5. Boxed region in the embryonic heart in (E) marking the endocardium and the myocardium are magnified in (E’) (N= 3 embryos for each genotype). White dotted line indicates the endocardium. Graph shows the quantitation of Smad2/3 level (N= 3 independent experiments). Statistical analysis was performed using one-sample t-test. Error bars indicate standard error of mean (SEM). *p<0.05, **p<0.01,***p<0.001, ****p<0.0001. Scale bar: (E) 100 µm; (E’) 20 μm.

**Fig. S3.**
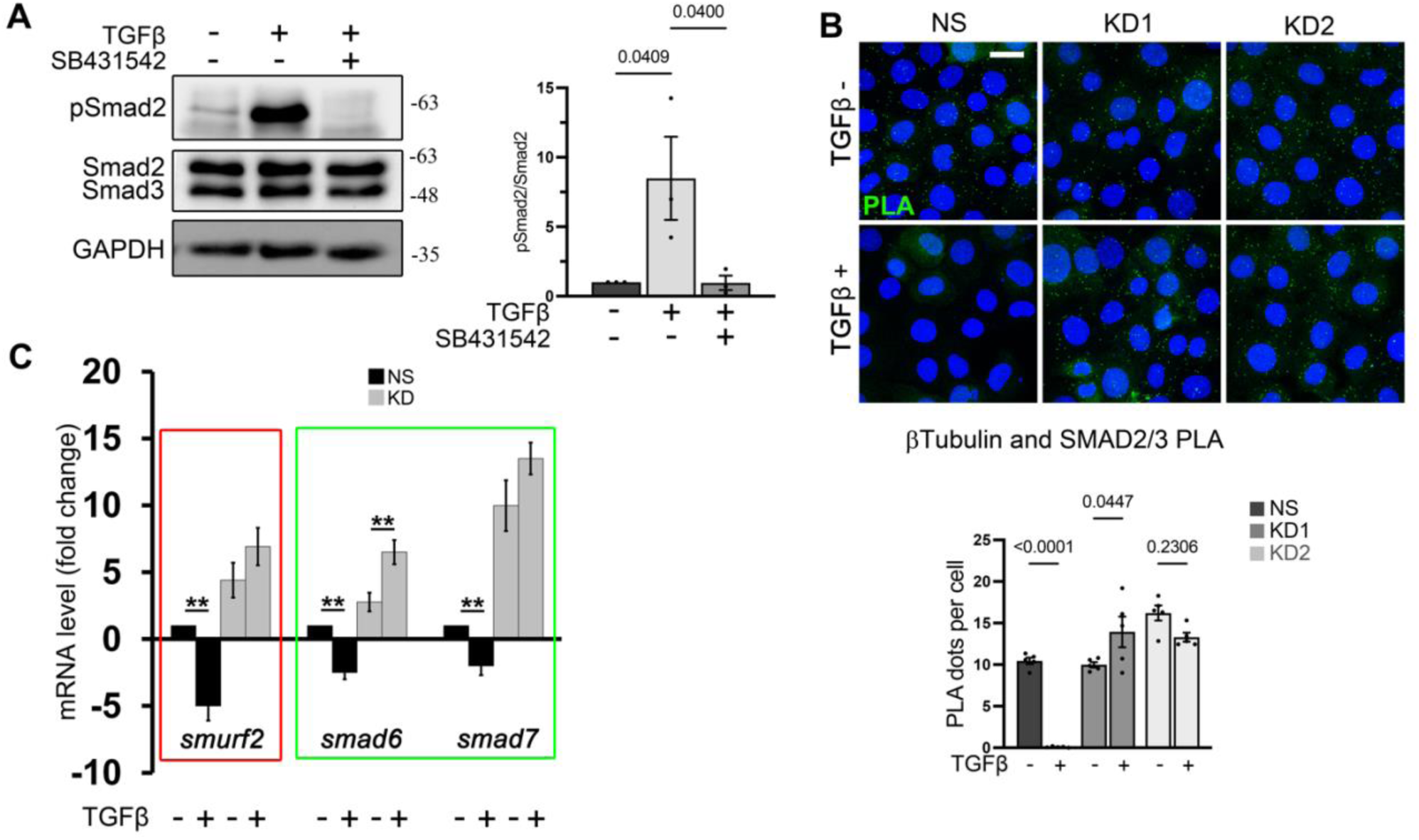
(related to Fig. 1, 2 and 3). Rudhira specifically regulates TGFβ-dependent Smad2/3 activation. (A) SVEC cells treated with SB431542 (10µM) for 2h and analyzed for Smad2 phosphorylation by Immunoblotting. (B) NS and two KD SVEC lines (KD1 and KD2) were analyzed for MT-SMAD interaction by PLA for β-Tubulin and Smad2/3 With or without TGFβ treatment (N= 3 independent experiments). (C) qRT-PCR analysis of negative regulators of TGFβ pathway (N=3 independent experiments). Statistical analysis was performed Using One way ANOVA. Error bars indicate standard error of mean (SEM). Scale bar (B): 20 μm. *p<0.05, **p<0.01, ***p<0.001.

**Fig. S4.**
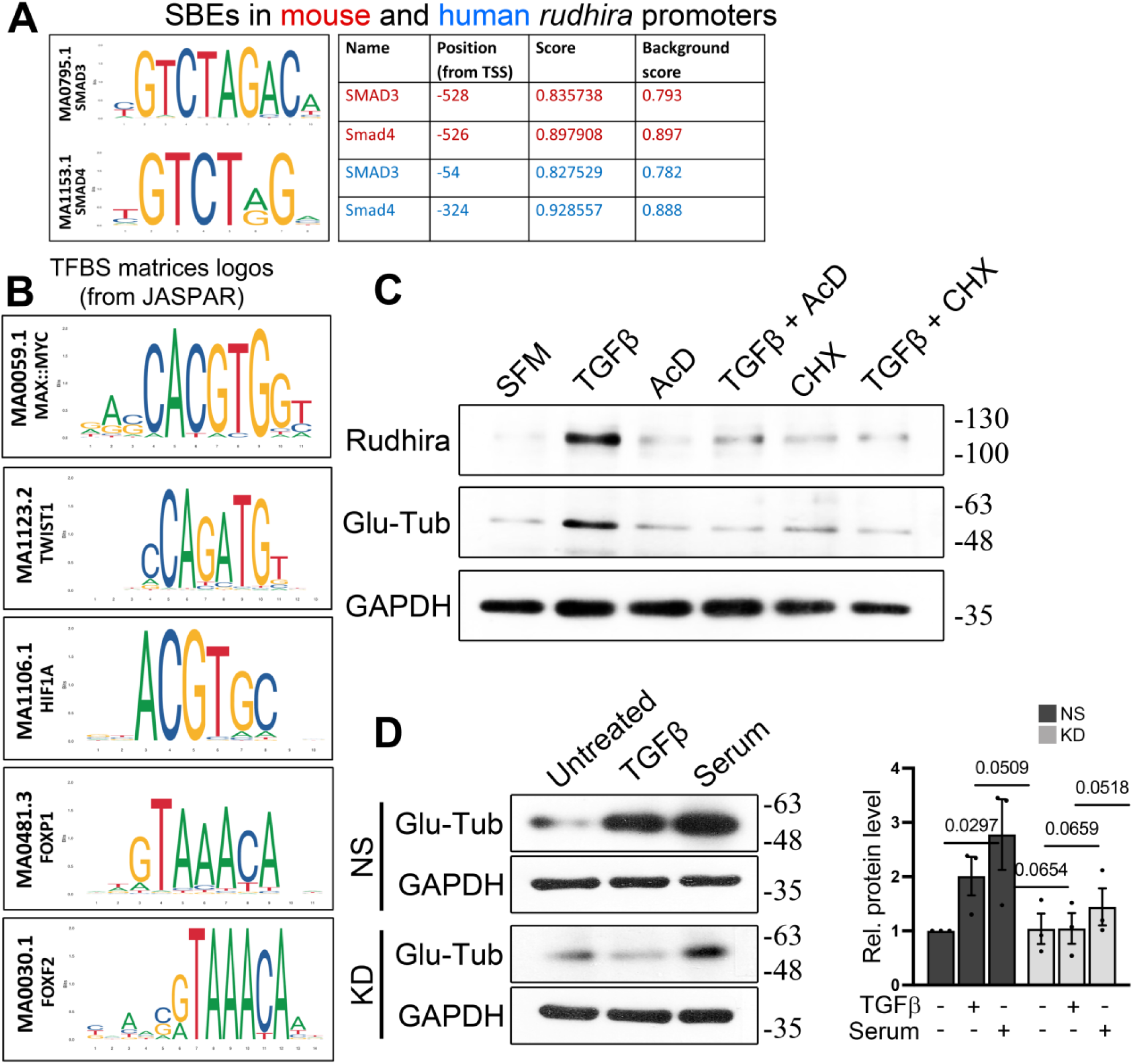
(related to Figs. 4 and 5). TGFβ induces *rudhira* transcription for MT stability. (A) Smad-binding element (SBE) matrix (SMAD3, ID MA0795.1; Smad4, ID MA1153.1) and predicted binding sites in mouse (in red) and human (in blue) *rudhira* promoters, from JASPAR and PSCAN bioinformatics tool. (B) Transcription factor binding sites (TFBS) available from JASPAR and PSCAN bioinformatics tool.(C) Analysis of Rudhira levels and MT stability (marked by Glu-Tubulin) upon treatments as indicated by immunoblotting (N= 3 independent experiments). (D) Analysis of MT stability in serum starved WT and KD cells kept untreated or treated with TGFβ or serum (N= 3 independent experiments).

## Supplementary Table S1.

**Supplementary Table S1.**
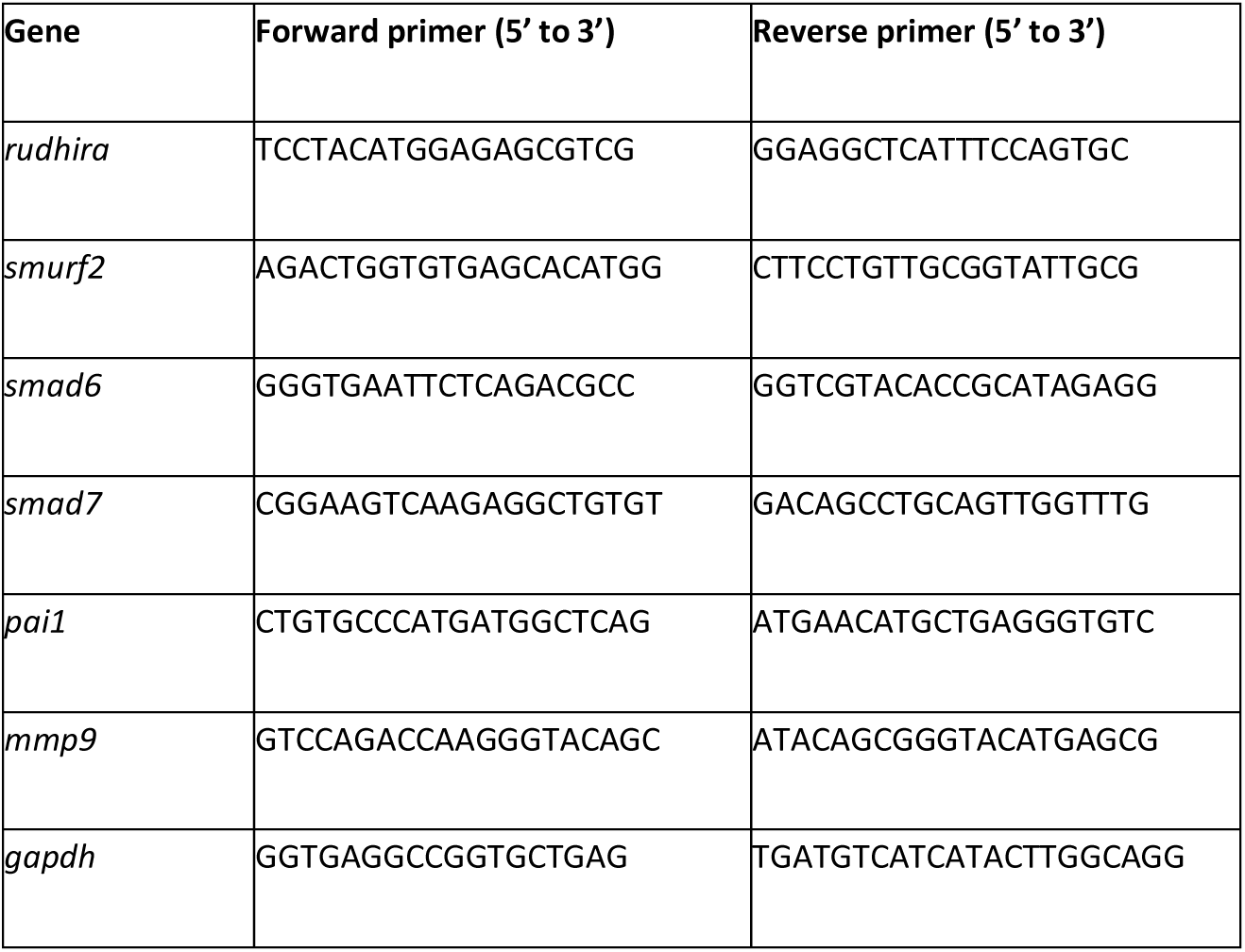
Primers used for qRT-PCR analysis.

**Supplementary Table S2.**
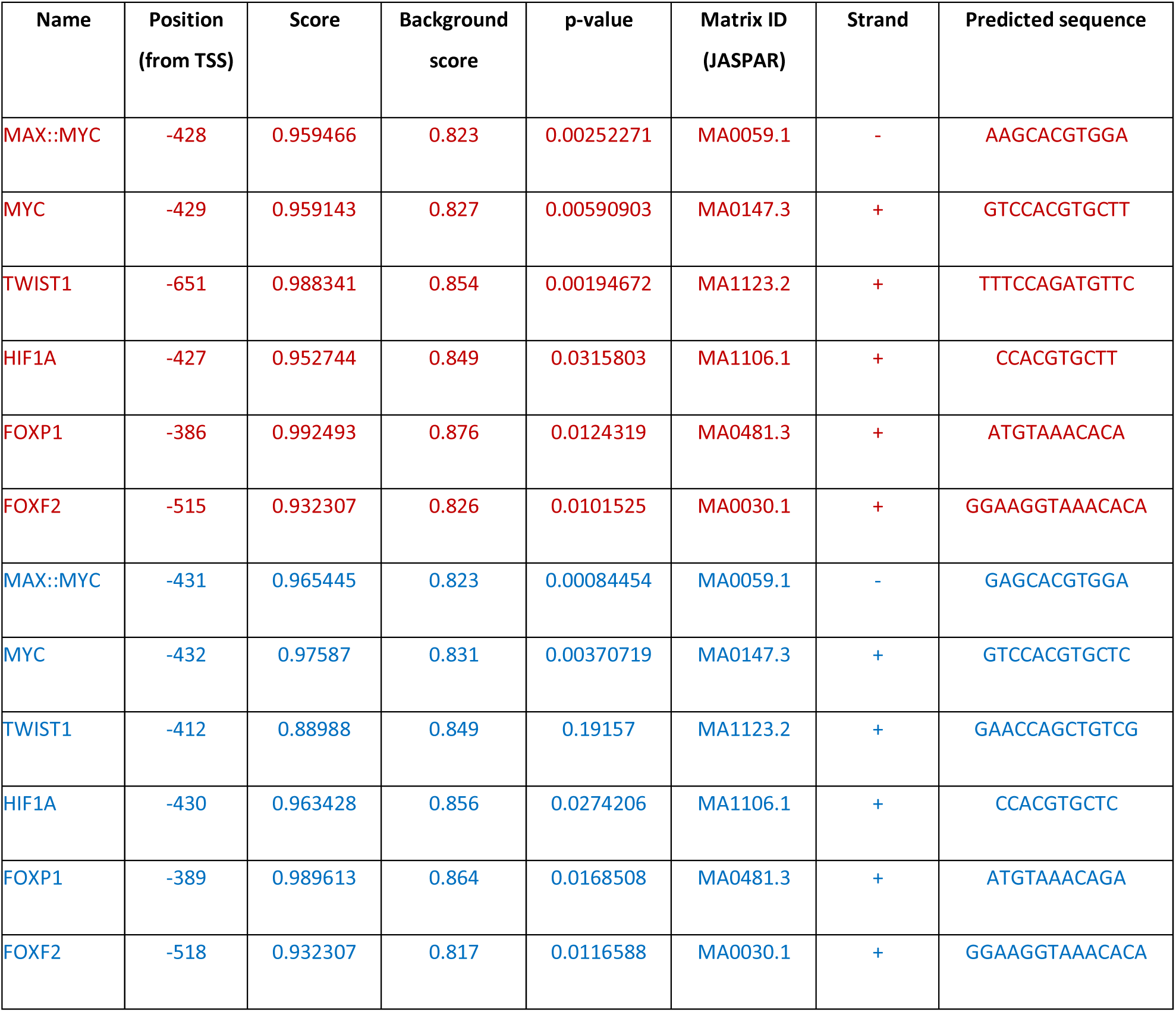
Predicted Transcription factor binding sites in mouse and human *rudhira/BCAS3* promoters, obtained from JASPAR bioinformatics tool.

**Supplementary Table S3.**
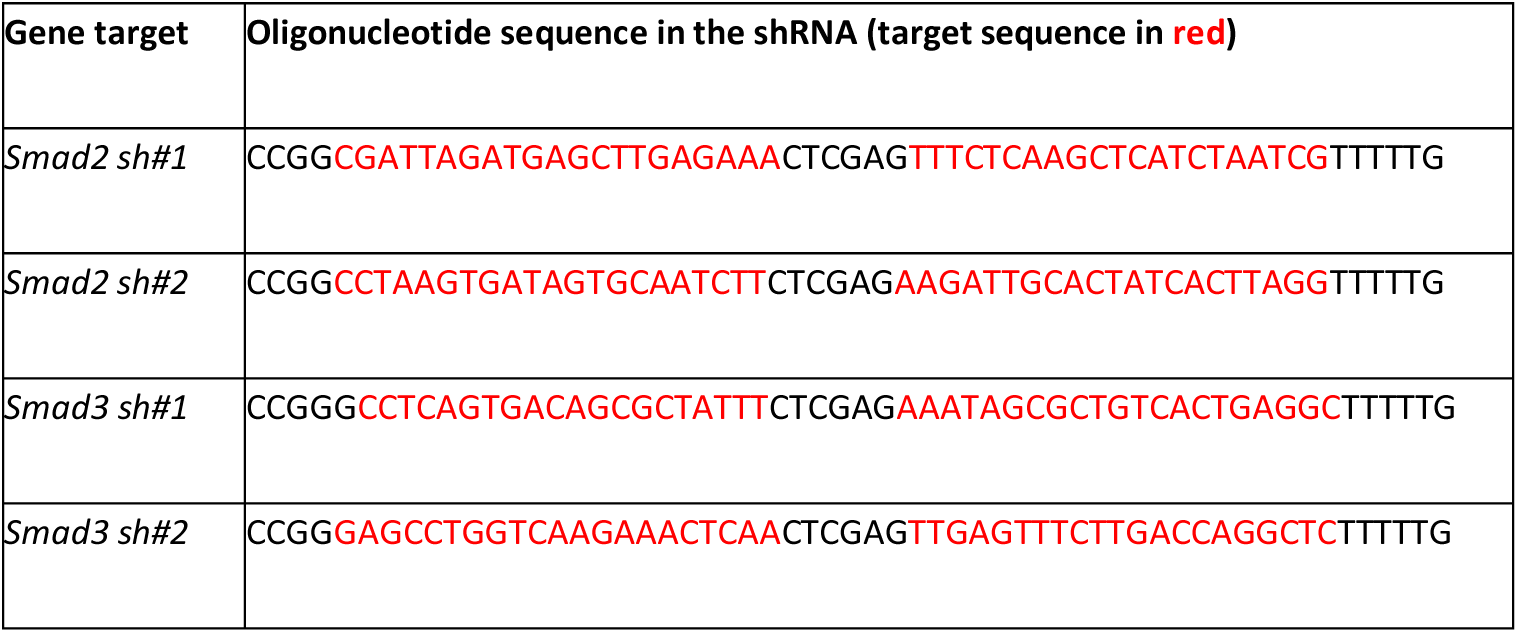
The oligonucleotide sequence of the *smad2* and *smad3* shRNAs.

